# Neural oscillations and top-down connectivity are modulated by object-scene congruency

**DOI:** 10.1101/2025.04.15.648958

**Authors:** Ye Gu, Alexandra Krugliak, Alex Clarke

## Abstract

The knowledge we have about how the world is structured is known to influence object recognition. One way this is demonstrated is through a congruency effect, where object recognition is faster and more accurate if items are presented in expected scene contexts. However, our understanding of the dynamic neural mechanisms that underlie congruency effects are under-explored. Using MEG, we examine how the congruency between an object and a prior scene results in changes in the oscillatory activity in the brain, which regions underpin this effect, and whether congruency results arise from top-down or bottom-up modulations of connectivity. We observed that prior scene information impacts the processing of visual objects in behaviour, neural activity and connectivity. Processing objects that were incongruent with the prior scene resulted in slower reaction times, increased low frequency activity in the ventral visual pathway, and increased top-down connectivity from the anterior temporal lobe and frontal cortex to the posterior ventral temporal cortex. Our results reveal that the recurrent dynamics within the ventral visual pathway are modulated by the prior knowledge imbued by our surrounding environment, suggesting that the way we recognise objects is fundamentally linked to their context.

---

The objects that we see and recognise are not experienced in isolation. It is well established that the process of object recognition is influenced by the scene an object is found within. This is demonstrated by faster and more accurate recognition rates for objects within congruent scenes compared to incongruent scenes (Bar, 2004; Davenport & Potter, 2004; Greene et al., 2015; Oliva & Torralba, 2007; Palmer, 1975), whilst visual scenes can also aid the disambiguation of ambiguous objects (Brandman & Peelen, 2017). However, we currently lack details about the neural underpinnings and mechanisms that are modulated by the relationship between an object and where it is located, which would give us a better understanding of object recognition under more naturalistic circumstances.

Object-scene congruency effects are typically explored in paradigms where an object is either embedded or overlaid on the visual scene, with most studies testing neural activity using EEG. Such research has widely found that neural responses are modulated approximately 250 ms after seeing the image, with various EEG components showing a sensitivity to object-scene congruency including the N300, N400 and P600, which have been linked to perceptual, semantic and structural processing of the object-scene relationship (Chen et al., 2022; Draschkow et al., 2018; Ganis & Kutas, 2003; Kumar et al., 2021; Lauer et al., 2018; Mudrik et al., 2014; Võ & Wolfe, 2013). While the modulation of ERPs is the dominant source of evidence for congruency effects, we have limited knowledge about how oscillatory processes might be modulated when it comes to object and scene congruence. Studies using congruency manipulations of word pairs or during sentence reading suggest that theta activity increases when there is a semantic violation or incongruency between the context and a new item (Bastiaansen et al., 2005; Packard et al., 2020). Increases in theta could reflect a modulation of retrieval processes during the semantic processing of the stimulus (Klimesch et al., 2001), with prior work also linking theta activity patterns to the semantic processing of objects (Clarke, 2020; Clarke et al., 2018). However, strikingly little research exists concerning the oscillatory nature of object-scene congruency effects.

A further issue is that we have little evidence for which neural regions are modulated by the recognition of objects in congruent or incongruent scenes. While N400 effects during reading and listening to words have been localised to language sensitive regions in the inferior frontal gyrus, posterior middle temporal gyrus, and the anterior temporal lobe (Ghosh Hajra et al., 2018; Halgren et al., 2002; Lau et al., 2008; Maess et al., 2006; Nobre & McCarthy, 1995), equivalent effects for object-scene congruency would be predicted in regions along the ventral temporal lobe that do display object-scene congruency modulations in fMRI (Li et al., 2023; Rémy et al., 2014) and are associated with the visual perception and integration of objects and scenes (Bar & Aminoff, 2003; Brandman & Peelen, 2017).

Going beyond this, an understanding of how scene knowledge influences object recognition processes requires that we test how connectivity is modulated, and whether top-down or bottom-up connections are impacted. While changes in activity due to congruency have been observed, such effects are limited in the ability to reveal the neural mechanisms that explain how those changes occur. If congruency results in modulations of regional activity, this could be due to differences in feedforward processes, top-down, or a combination. Assessing how network connectivity is altered, and which regions drive these changes, is essential to gain a fuller understanding of how a preceding scene context changes the neural mechanisms underpinning the processing of visual objects.

We explored these issues using MEG, where objects were preceded by a visual scene that was either congruent or not with the object. Through analysing MEG sensor activity, time frequency responses, source localised activity, and effective connectivity, we asked if the prior scene context influenced the processing of visual objects through changes in oscillatory activity, which neural regions underpin this effect, and finally, how connectivity dynamics are modulated within this neural system by congruency.

## Materials and Methods

### Participants

Thirty-one participants (12 males, age range 18-35 years) took part in the study. All participants were right-handed and had normal or corrected-to-normal vision. Three participants were excluded due to poor behavioural performance (object recognition accuracy less than 65%) leading to a final sample size of 28. All participants gave written informed consent. The study was approved by the Cambridge University Psychology Research Ethics Committee (PRE.2019.051).

Stimuli A total of 450 colour images were used in the study, including 150 scene images, 150 scrambled scene images, and 150 object images. The scenes images were obtained from Lauer et al. (2018) the SUN397 scene image database (Xiao et al., 2010), and internet searches using Google Image Search. Objects were shown in colour, isolated on a white background, from the Hemera photo object image set or Clarke et al. (2018).

Each scene image was paired with two object images, one that was congruent with the scene, and one that was incongruent. Objects only appeared once with a congruent scene and once with an incongruent scene, meaning that each scene was paired with two objects, and each object was paired with two scenes. The object-scene pairings were determined with a pretest using a separate group of 37 participants and a larger range of scene and object images. The pairings were initially composed by the research team prior to the pretest. Participants rated each object-scene pairing based on how likely they were to encounter the object within the scene using a 5-point scale (1 = very unlikely, 2 = unlikely, 3 = neutral, 4 = likely, 5 = very likely). Congruent pairings were defined as those with a mean rating above 3.5 and incongruent pairings with a mean rating of less than 1.5. If a pairing did not have a clear congruent or incongruent rating, the images were adjusted and rated again, resulting in 150 congruent pairs and 150 incongruent pairs where each scene and object appeared once in each list. Importantly, the objects did not appear in the scenes they were paired with.

The scrambled scene image that was paired with each object was created from the two scene images paired with that object. The two scene images were divided into small squares and a new image was created by combining half the squares from one scene with half from the other scene. The squares were flipped and randomly shuffled to create a scrambled image. The condition with scrambled images is not analysed in this study.

### Paradigm

Participants performed an object recognition task, where an object image followed a scene that was either congruent or incongruent with the object, or a scrambled scene image. Each trial began with a fixation cross lasting 750 ms before a scene image was shown for 500 ms. This was followed by a blank grey screen lasting 1 second. An object was then displayed in the centre of the screen for 50 ms followed by a backwards mask lasting 200 ms. The mask was made of dynamically changing overlapping squares with each pattern lasting 50 ms (Figure 1a).

**Figure 1.**
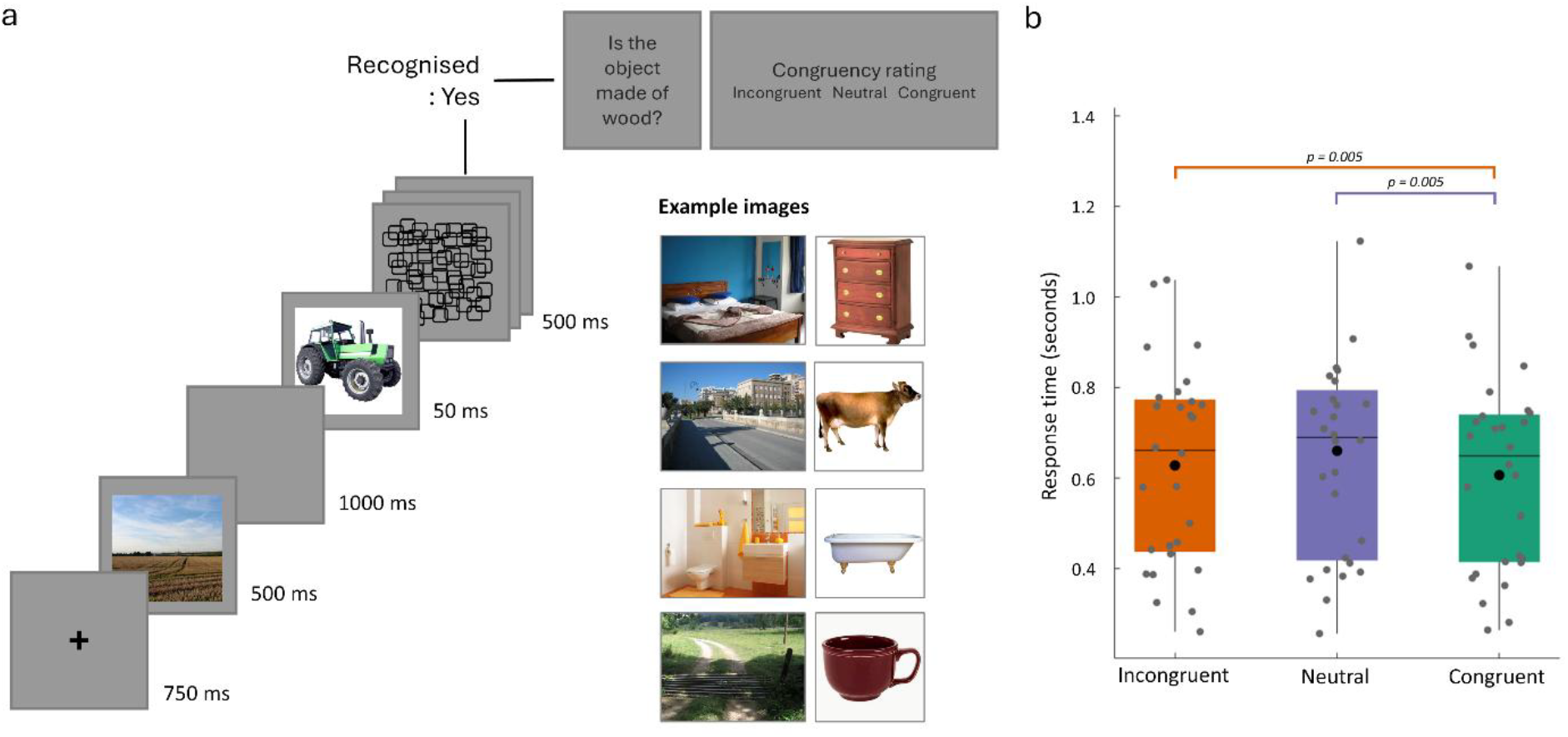
Experimental paradigm and behavioural performance. a) On each trial, a scene is presented followed by a backward masked visual object. Participants make a button press if they saw and knew what the object was. A catch question is used to ensure they recognised the object, and participants are then asked to rate if the prior scene was congruent with the object on a 3-point scale. Example images show the range of congruencies between the scenes and the objects. b) Behavioural response times for recognised objects, classified according to the congruency ratings. Congruent object-scene trials were significantly faster than both incongruent and neutral rated trials.

Participants pressed a button using their right index finger if they recognized the object and knew what the object was (not just that they saw something), or using their right middle finger if they did not recognize the object. For objects that were recognized, they next answered a yes/no question about the object’s characteristics (e.g., ‘‘Is the object edible?’’, or ‘‘Can you buy the object in a supermarket?’’), followed by a congruency rating for the object and scene pair. Congruency ratings were made using the left hand and with 3 response options (unlikely, neutral, likely) to the question ‘How likely would you expect to encounter this object in this context?’.

If the participant did not recognize the object, no following questions were asked. Each object was shown three times, on each occasion being preceded by the congruent, incongruent or scrambled scene image, with a total of 450 trials. Each participant saw the 450 trials in a randomized order. The experiment was presented on a screen positioned 120 cm from the participant with a resolution of 1024×768 (approximately 57×33 cm). All scene images were full screen with a resolution of 1024×768, and object images had a resolution of 500×500 pixels. The experiment was controlled using MATLAB (MathWorks, Natick, MA) and Psychtoolbox (Brainard, 1997).

### MEG/MRI acquisition

Continuous MEG data were recorded using a whole-head 306 channel (102 magnetometers, 204 planar gradiometers) TRIUX neo system (MEGIN, Espoo, Finland) at the MRC Cognition and Brain Sciences Unit, Cambridge. Five head-position indicator (HPI) coils were used to record the head position (every 200 ms) within the MEG helmet and eye movements and blinks were monitored using the electrooculogram (EOG) recorded bipolarly through electrodes placed above and below the right eye (vertical) and at the outer canthi (horizontal). The participants’ head shape was digitally recorded using a 3D digitizer (Fastrak Polhemus, Inc., Colchester, VA, USA) along with the positions of the EOG electrodes, HPI coils, and fiducial points (nasion, left and right periarticular). MEG signals were recorded at a sampling rate of 1000 Hz, with a high-pass filter of 0.03 Hz.

To facilitate source reconstruction, high-resolution structural T1-weighted MPRAGE scans were acquired during a separate session with a Siemens 3T Tim Trio scanner (Siemens Medical Solutions, Camberley, UK) located at the MRC Cognition and Brain Sciences Unit, using a 3D MPRAGE sequence (field-of-view 256 mm × 240 mm × 160 mm, matrix dimensions 256 × 240 × 160, 1 mm isotropic resolution, TR = 2250 msec, TI = 900 msec, TE = 2.99 msec, flip angle 9°).

### MEG processing

The raw data were Maxfiltered using MNE-python (https://mne.tools/). Static bad channels were detected and subsequently reconstructed by interpolating neighbouring channels, as were bad channels containing long periods of high amplitude or noisy signals. The temporal extension of the signal-space separation (SSS) technique (Taulu et al., 2005) was applied to the data every 4s to segregate the signals originating from within the participants’ heads from those generated by external sources of noise, along with head movement compensation and transformation of the head position to a common position.

The resulting MEG data were processed using SPM12 (Wellcome Institute of Imaging Neuroscience, London, UK, www.fil.ion.ucl.ac.uk/spm), and were downsampled to 250 Hz, low-pass filtered at 100 Hz, stop-pass filtered between 48 and 52 Hz, and high-pass filtered at 0.1 Hz in forward and reverse directions using a fifth-order bidirectional Butterworth digital filter. The data were then epoched from -2500 to 2000 ms relative to the onset of the object (or -1000 to 3500 ms relative to the onset of the scene). Independent components analysis (ICA) was implemented using EEGLab (Delorme & Makeig, 2004), and components whose time series showed a Pearson’s correlation greater than 0.4 with horizontal or vertical EOG channels were labelled for rejection. Additional manual inspections identified components involving apparent rhythmic cardiac noise, before MEG signals were reconstructed using the montage function to retain the non-identified components. Epoch were baseline corrected using the 200 ms period prior to the object, and trials that were not recognized were rejected (mean = 18, range 0 - 74 trials, SD = 19.2 trials).

Our statistical analysis of the data used Fieldtrip (Oostenveld et al., 2010). The SPM files were converted into fieldtrip format (spm2fieldtrip.m), trials were re-labelled according to the participant ratings and event-related averages calculated for each participant. Our main analyses only used the congruent and incongruent trials as there were fewer trials rated as neutral (congruent: mean = 118, range 89-146 trials; incongruent: mean = 123, range 72-155 trials; neutral: mean = 39, range 7-127 trials). To test for congruency effects across sensors and time-points, we used a cluster-based permutation test, focusing on time points between 0 and 1250 ms with a cluster-forming alpha of 0.01, 10,000 randomisations, and a minimum number of cluster-forming channels set to 2. The analysis time window was chosen to encompass the object onset and previous congruency modulations of EEG activity seen for N300, N400 and P600 components (e.g. Võ & Wolfe, 2013). We also used an analysis of time frequency representations to test for congruency effects. Time frequency representations were calculated for each trial using Morlet wavelets of length 5 cycles at 40 logarithmically spaced frequencies between 2 and 100 Hz. A baseline correction was applied to convert the data into dB using data between -500 and -200 ms prior to the onset of the object, and the data was averaged across trials. To test for congruency effects across sensors, frequencies and time-points, we used a cluster-based permutation test, focusing on time points between 0 and 1250 ms and frequencies between 3 and 40 Hz. We used a cluster-forming alpha of 0.01, 10,000 randomisations, and a minimum number of cluster-forming channels set to 2.

We also performed additional analyses within clusters identified by the contrast between congruent and incongruent trials to see if the effects further distinguished between the three possible levels of congruency (congruent, neutral and incongruent). By using the mean amplitude across a time window and selection of sensors, we can compare conditions with large differences in the number of trials per condition (Luck, 2014). Repeated measures ANOVA and follow-up tests were run using JASP 0.18.3.0. Greenhouse-Giesser correction to the degrees of freedom was used for violations of sphericity, and multiple follow-up t-tests were corrected for multiple comparisons using the Holm-Bonferroni correction.

### Source localization

To localize the time frequency effects we used a beamforming approach in SPM12. Individual participants MRI images were segmented and spatially normalised to an MNI template brain consisting of 20,484 vertices, which was inverse normalised to the individuals specific MRI space. MEG sensors were co-registered to the MRI image using the three fiducial points and the additional headpoints. A single shell forward model was used. A linearly constrained minimum variance (LCMV) beamformer was implemented using the DaiSS toolbox. Only the magnetometers were included in the inversion, with a frequency range set between 6 and 12 Hz, a time window of 200 to 1000 ms, and a grid size of 5 mm. Voxel-based images showing the source activity for each participant and the two conditions of interest, congruent and incongruent, were created in MNI space, and smoothed using a 10 mm FWHM kernel. Source activity values were log transformed. These images were entered into a paired-samples T-test, assessed using a voxel-wise threshold of p < 0.01, and a cluster threshold of p(FWE) < 0.05. Images are displayed on an inflated surface using BrainNet viewer (Xia et al., 2013).

### Effective connectivity

We used dynamic causal modelling (DCM) for cross-spectral density (Friston et al., 2012) to assess the directed effective connectivity between regions identified in the beamformer analysis outlined above. Using the peak coordinates from the beamformer analysis, we specified a two-node network consisting of the left posterior temporal cortex (MNI coordinate -18, -50, -6) and left anterior temporal lobe (MNI coordinate –32, -10, -34). An additional three-node network also included the left inferior frontal gyrus identified from a meta-analysis of semantic control (MNI coordinate -42 34 -6; Noonan et al., 2013). For both networks, the connections set out by the A matrix included bi-direction connections, which were modulated in the B matrix. The frequency range was set to 6-12 Hz, and the time window between 0 and 1000 ms after the object appeared, with the congruent condition treated as a baseline. To make inferences about how feedforward or feedback connectivity was modulated by congruency, we assessed the group-level modulations using the PEB framework, using Bayesian model reduction (BMR) and Bayesian model averaging (BMA) (Zeidman et al., 2019).

### Behavioural stats

For each participant, we collected object recognition reaction times and the rated congruency between the object and the preceding scene. Reaction times for successful recognition trials were classified into congruent, incongruent and neutral conditions based on the participants own ratings. Repeated measures ANOVA and follow-up tests were run using JASP 0.18.3.0. Greenhouse-Giesser correction to the degrees of freedom was used for violations of sphericity, and multiple follow-up t-tests were corrected for multiple comparisons using the Holm-Bonferroni correction.

## Results

### Behaviour

Analysis of the behavioural responses show that participants were able to recognize the briefly presented objects with a high degree of accuracy (mean = 96%, minimum 79%, maximum 100%). From the trials that were successfully recognised, we tested for an effect of congruency on response times using the condition labels given by individual participants (Figure 1b). A repeated measures ANOVA with factor congruency (congruent, neutral, incongruent) showed a significant effect of congruency (F(1.18,31.8)=7.4, p=0.008). Posthoc comparisons showed that congruent trials were responded to fastest compared to both incongruent (t(27)=3.44, p=0.005) and neutral trials (t(27)=3.48, p=0.005), while no significant differences were observed between incongruent and neutral trials (t(27)=1.82, p=0.08). These results align with past research where objects are recognized faster when they are related to a congruent scene (Bar, 2004; Davenport & Potter, 2004; Greene et al., 2015; Oliva & Torralba, 2007; Palmer, 1975).

**Table 1.**
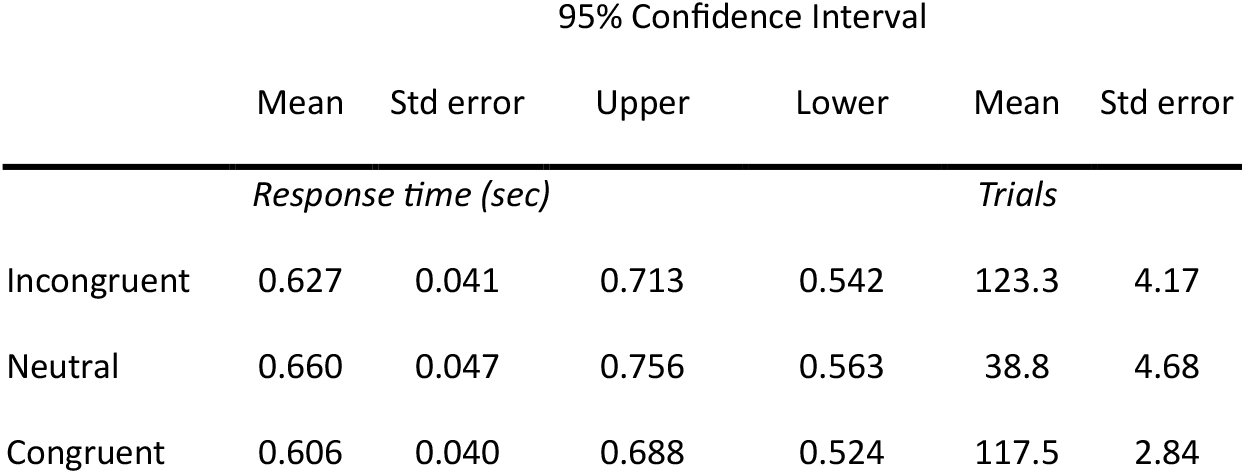
Behaviour performance.

### MEG sensor-level analysis

Before testing our main research question, we first established that our manipulation of object-scene congruency evoked effects similar to N300/N400 effects widely seen in EEG studies, despite our temporal separation of the scene from the object. To ensure the effects related to each individual participants view of congruency, we created conditions based on the subjective judgements of each participant.

Our analysis focused on the time period after the object was presented, testing for differences between the congruent and incongruent trials across MEG sensors and time points. A cluster-based permutation test revealed a significant effect where incongruent trials showed larger negative field strength compared to congruent trials over the left anterior sensors between approximately 500 and 700 ms after object onset (cluster p = 0.0349; Figure 2a,b,c). Whilst the significant cluster might look slightly later than many N300/N400 ERP effects, examination of the time-series suggests the effect might be temporally more extended, from around 200 to 1000 ms (Figure 2b). To further explore how this effect relates to a fuller range of congruency, the mean amplitude from this cluster was compared across the congruent, neutral and incongruent trials (Figure 2c). A repeated measures ANOVA with factor congruency (congruent, neutral, incongruent) showed a significant effect of congruency (F(1.5,39.7)=8.075, p=0.003). Posthoc comparisons showed that congruent trials had a greater mean amplitude compared to both incongruent (t(27)=4.42, p<0.001) and neutral trials (t(27)=3.41, p=0.004), while no significant differences were observed between incongruent and neutral trials (t(27)=0.98, p=0.4). This shows that our paradigm and data elicit congruency effects during the processing and integration of objects with the prior scene.

**Figure 2.**
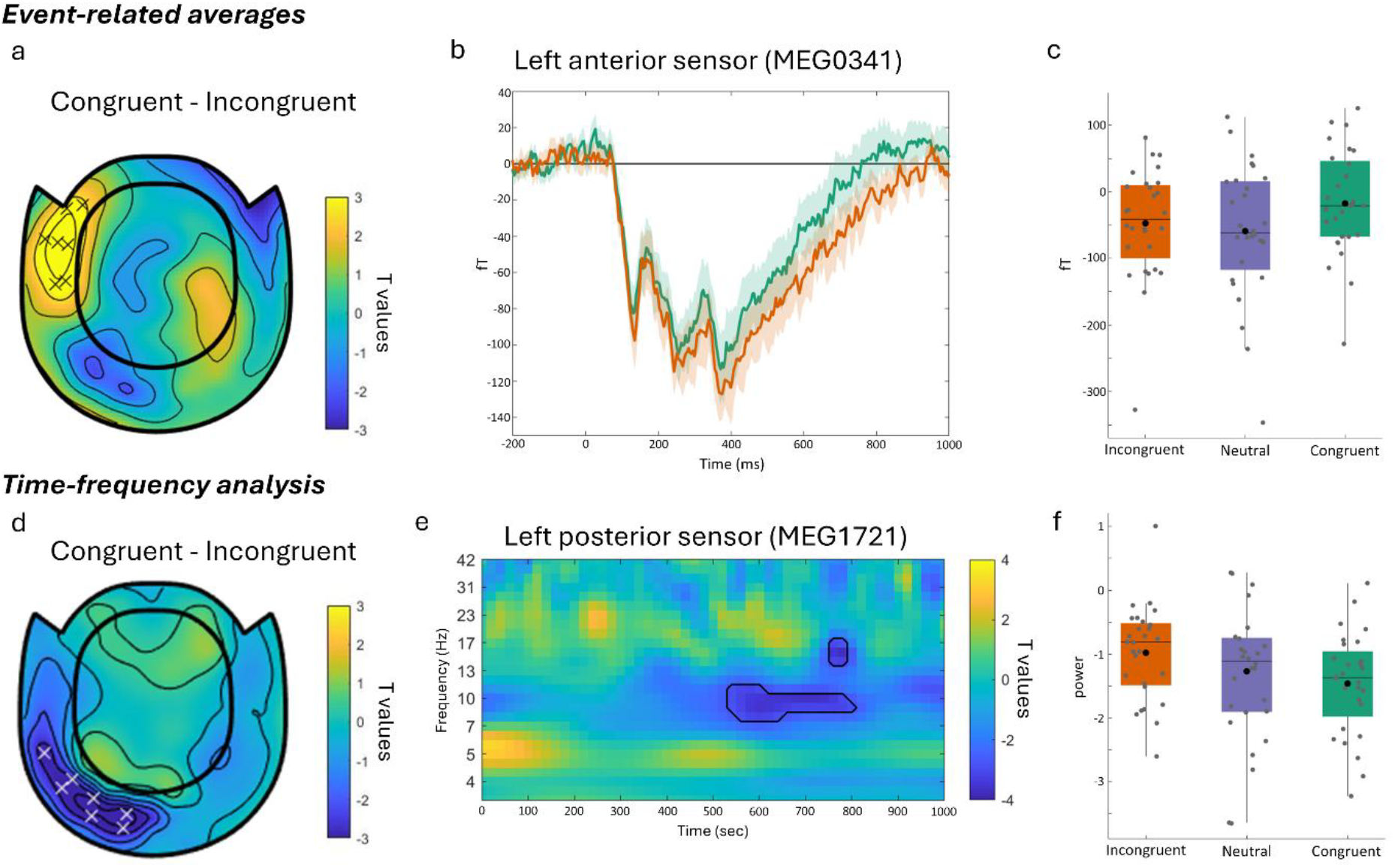
Sensor-level congruency effects. a) Event-related average analysis showing a representative topography of the congruency effect. Data shows effect at 600 ms, with ‘x’ indicating sensors showing a significant effect. b) Event-related activity at a representative sensor for the congruent and incongruent conditions. c) Data averaged across time-points and sensors from the significant cluster, plotted for the congruent, incongruent and neutral trials. Each data-point is one participant. d) Time-frequency analysis showing representative topography of the congruency effect, plotted at 8.6 Hz and 560 ms. ‘x’ indicates sensors showing a significant effect. e) Time frequency representation for a representative posterior sensor showing t-statistics for the contrast of congruent – incongruent trials. Black line shows the extent of the significant cluster at this sensor. f) Data averaged across time-points, sensors and frequencies from within the significant cluster, plotted for the congruent, incongruent and neutral trials. Each data-point is one participant.

One of our main research questions was how neural oscillations are modulated by object-scene congruency. Whilst modulations of activity seen though ERPs and ERFs have been observed, far less attention has been paid to modulations of oscillations. To address this, we calculated the power at each frequency between 3 and 40 Hz and for time points between 0 and 1250 ms for each trial and at every MEG sensor. This allowed us to test if total power at different frequencies was modulated by the congruency between the object and prior scene, and over which sensors any effect was located.

We used a cluster-based permutation test for an effect of congruency across MEG sensors, time and frequency. This showed a significant modulation of low frequency activity, where more power was seen for incongruent trials compared to congruent trials, with the effect peaking around 8 Hz, over left posterior sensors, and between approximately 500 to 900 ms after object onset (cluster p = 0.0304; Figure 2d,e,f). Again, this appears part of a wider divergence between the conditions from near 200 ms to over 1 second, and across frequencies near 6-12 Hz in the high theta and alpha range (Figure 2f).

To further explore this effect across the range of congruency responses, the mean amplitude from this cluster was extracted (Figure 2f). A repeated measures ANOVA with factor congruency (congruent, neutral, incongruent) showed a significant effect of congruency (F(1.3,35.3)=5.37, p=0.019). However, posthoc comparisons only show the difference between congruent and incongruent trials (t(27)=6.16, p<0.001), with no significant differences between the neutral trials and congruent (t(27)=1.08, p=0.29) or incongruent trials (t(27)=1.74, p=0.19).

### Neural network modulation by congruency

In order to understand which neural regions were underpinning the low frequency modulation by congruency, we performed a source localisation procedure incorporating the individuals anatomical MRI data. We used an LCMV beamformer to determine how activity was distributed across the brain for the congruent and incongruent conditions for each participant, with activity estimated for the time window between 200 and 1000 ms, and between 6 to 12 Hz. This revealed that objects that are incongruent with the preceding scene were associated with significantly increased low frequency activity in the left ventral visual pathway, from the posterior ventral temporal cortex (VTC) through to the anterior temporal lobe (ATL) (voxelwise p < 0.01, cluster p < 0.05; Figure 3).

**Figure 3.**
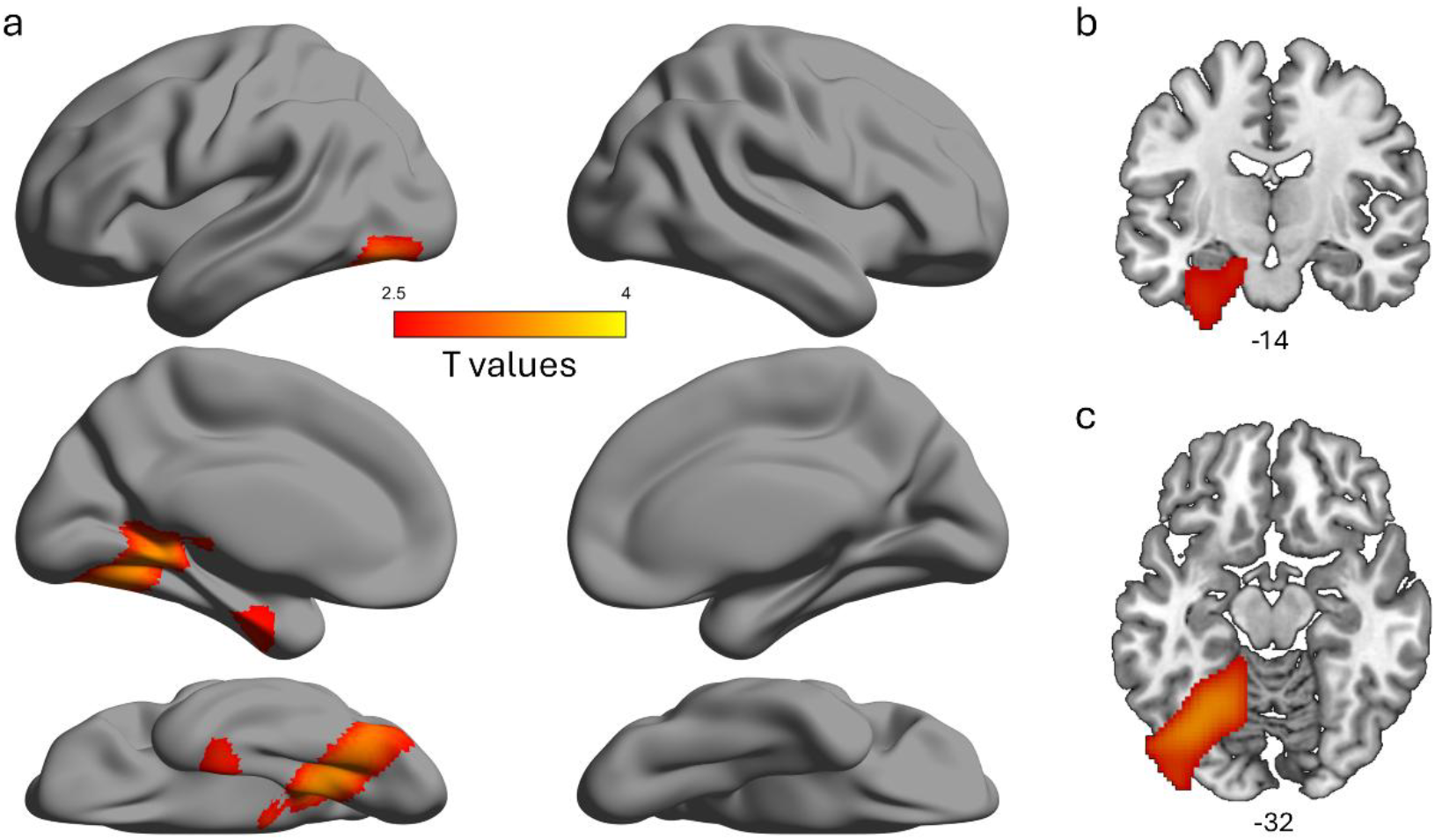
Source localisation of the low frequency congruency effect showing increased activity for the incongruent condition compared to the congruent condition, shown a) on an inflated surface, b) coronal view and c) axial view. Thresholded at p < 0.01, cluster p < 0.05.

Finally, we asked if connectivity in this network across the anterior and posterior ventral temporal lobe was modulated by congruency. We used DCM of cross-spectral densities to determine the most likely directional-network model that explains our low-frequency modulations according to congruency. Our network consisted of two regions in the left ventral visual pathway, defined by peaks in our source analysis above – the left VTC and left ATL. We defined this network as having feedforward and feedback anatomical connections between them, and we tested if these were modulated by congruency or not.

We used the second-level PEB framework to assess the probability that the feedforward or feedback connection was modulated by congruency. After estimating the connectivity parameters of the full model, we compared the full and reduced models using Bayesian Model Comparison and Reduction, where each of the parameters is switched on or off to determine the best network model of the data. This showed that the model that included a feedback modulatory connection had the highest posterior probability (83%) with low probability for models with either feedforward modulations only, or both feedforward and feedback modulations (Figure 4a).

**Figure 4.**
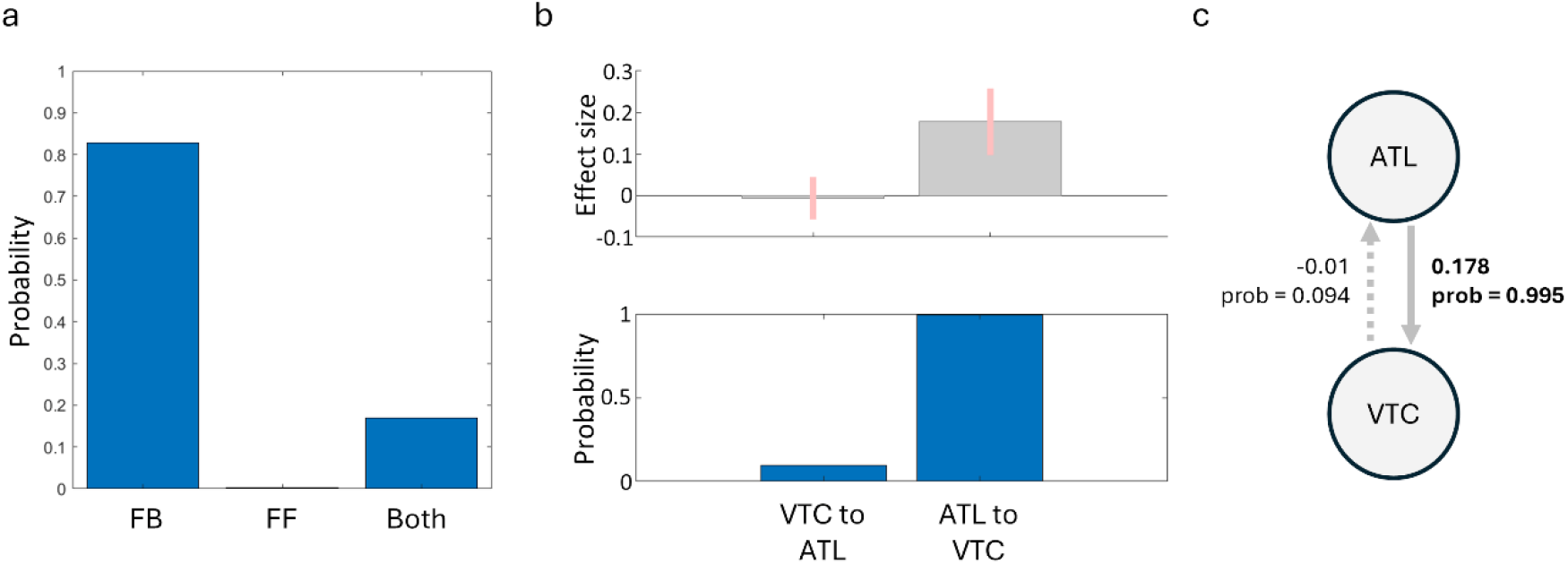
Congruency modulates feedback connectivity in the ventral visual pathway. a) Model probabilities when a given modulatory parameter is included. b) Bayesian model average parameters weighted by the model probabilities, showing the modulatory effect of the feedforward and feedback connection in explaining the congruency effect. Error bars show 95% confidence intervals. c) Summary of the modulatory effect of congruency, showing increased feedback from the ATL for the incongruent condition compared to the (baseline) congruent condition. Bold connections have probability greater than 99%.

To better understand the parameters, we performed Bayesian Model Averaging across the three model-types, where the parameters are weighted according to the probabilities of the model. This again indicates that while there is little evidence that feedforward connections were modulated by congruency (9.4% probability), there was strong evidence that feedback connectivity was increased for the incongruent condition compared to the congruent condition (99.5% probability; Figure 4b,c). This shows that when objects are incongruent with the prior scene, there is an increase in feedback connectivity from the ATL to the VTC (Figure 4c).

In addition to our 2-node analysis, we additionally investigated a 3-node network given the role the LIFG has during semantic control processes (Jefferies, 2013; Lambon Ralph et al., 2017). Like above, we tested which of the feedforward and feedback connections were modulated by congruency or not. After estimating the parameters for the full model, model probabilities were calculated for each of the possible 64 model permutations where each combination of modulatory parameters are switched on or off. To determine the nature of the model that best explains the data, probabilities were calculated for different families of the network. This shows that a network model with modulations of feedback connections has the highest probability (89.7%), and that a network model with modulations that involves all three nodes has the highest probability (83.7%). When comparing all possible models, while one model had clearly the highest probability (model 11, 47.4%), no single model could be described as an overall winner (i.e. probability >95%).

To better understand the modulatory effects in this network, we performed Bayesian Model Averaging across all models with the parameter estimates weighted by the model probabilities. In agreement with our 2-node analysis, this shows that there is strong evidence that feedback connectivity from the ATL to the VTC was increased for the incongruent condition compared to the congruent condition (99.7% probability; Figure 5d,e). The 3-node analyses additionally found strong evidence that feedback connectivity from the IFG to the VTC was increased for the incongruent condition compared to the congruent condition (98.6% probability; Figure 5d,e). This shows that when objects are incongruent with the prior scene, there is not only increases of within-temporal feedback, but that feedback from frontal regions also increases.

**Figure 5.**
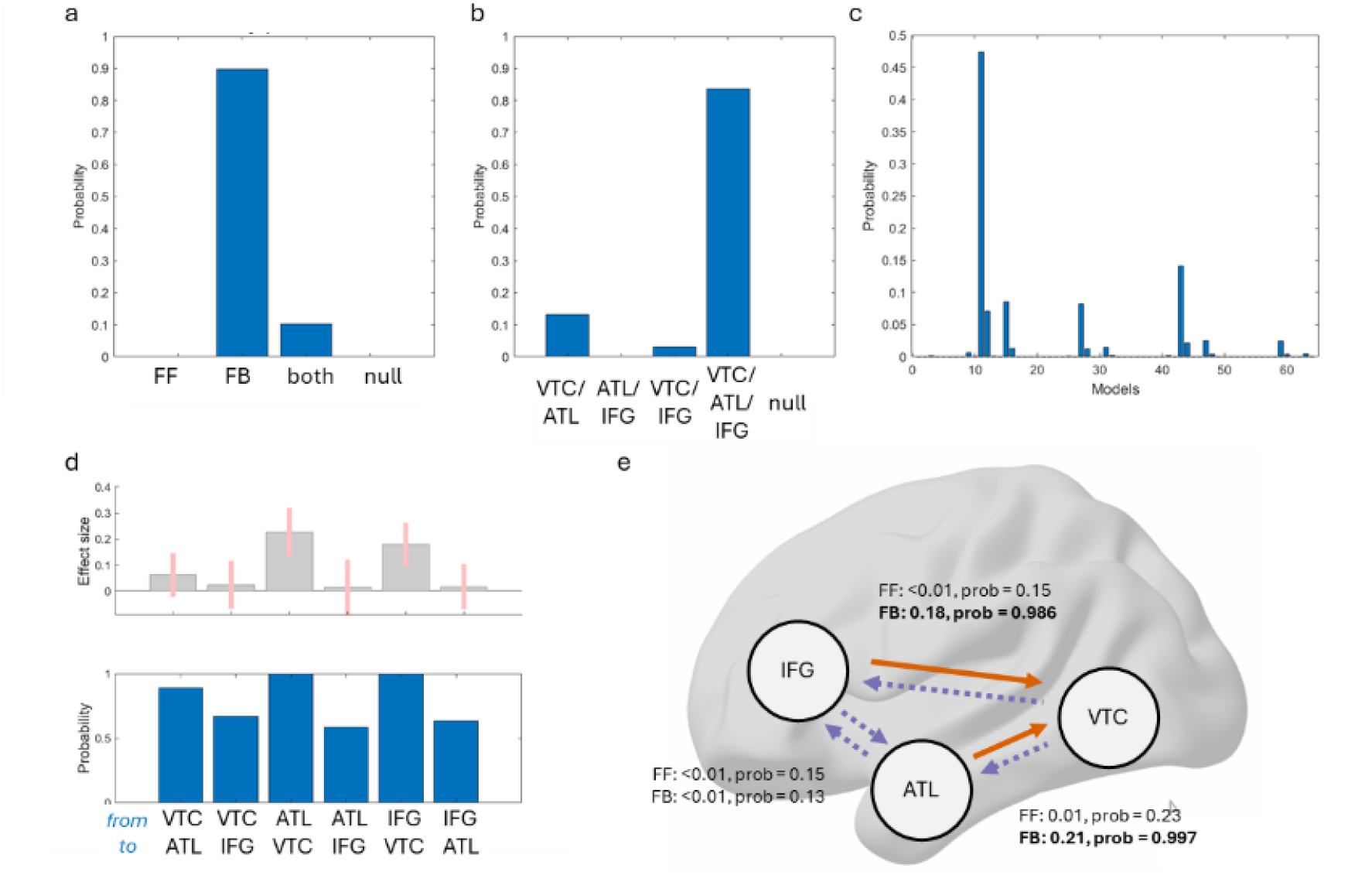
Congruency modulates feedback connectivity to the ventral temporal cortex. a) Model probabilities when different combinations of (a) feedforward and feedback connections are included, and (b) connections are included. c) Model probabilities of the entire model space tested, where each connectivity combination is selectively modulated. Model 11 shows the largest evidence, which specifies modulatory feedback from the IFG to the VTC, and from the ATL to the VTC. d) Bayesian model average parameters weighted by the model probabilities in (c), showing the modulatory effect sizes and probabilities of the feedforward and feedback connections in explaining the congruency effect. Error bars show 95% confidence intervals. e) Summary of the modulatory effect of congruency from the Bayesian model averaging, showing increased feedback from the ATL and from the IFG to the VTC for the incongruent condition compared to the (baseline) congruent condition. Bold/orange connections have probability greater than 98%.

## Discussion

We aimed to reveal how prior knowledge, established through a visual scene context, influenced subsequent object processing. Using MEG, we tested whether scene-object congruency impacted oscillatory activity, where this activity localised to in the brain, and if the connectivity dynamics of this system were modulated in a top-down or bottom-up fashion. We observed that prior scene information clearly impacts the processing of visual objects through behaviour, neural activity and connectivity, finding that processing objects that are incongruent with the prior scene leads to slowed reaction times, an increase in low frequency activity in the ventral visual pathway, and an increase in top-down connectivity. These results suggest that the neural processing and mechanisms underlying object recognition do not occur in isolation, but are sensitive to the environment visual objects are experienced within.

While models of object recognition are increasingly incorporating information about temporal dynamics, other research shows that the prior cognitive state of the brain influences subsequent visual processing (Aitken et al., 2020; de Lange et al., 2013; Dehaghani & Zarei, 2025; Mayer et al., 2016). For instance, when an auditory cue is predictive of an upcoming visual stimulus, the early sensory processing of that stimulus is modulated by the prediction cue (Aitken et al., 2020). In our paradigm, any predictions or expectations generated by the initial visual scene could lead to modulations of object processing, as seen through congruency between the scene and the object. We observed that top-down connectivity to the VTC increased when the object was incongruent with the prior scene. Using DCM for cross-spectral densities, we observed that a model where feedback connectivity increased for incongruent compared to congruent conditions was the best explanation for the observed changes in low frequency activity in the ventral visual pathway. Top-down connectivity to the VTC during object recognition has been observed under challenging recognition conditions such as visual occlusion, degraded images and manipulations of consciousness (Bar et al., 2006; Ganis et al., 2007; Mohsenzadeh et al., 2018; Rajaei et al., 2019; Schendan & Ganis, 2015; Schendan & Stern, 2008; Wyatte et al., 2014). Our results further show that our prior expectations about what object might appear also modulates top-down effects, and we suggest that this reflects the semantic processing of the objects which either do, or do not benefit from prior scene information.

Our connectivity analysis identified contextually modulated top-down effects from the IFG and ATL. The LIFG is part of a semantic control network, which supports the flexible retrieval of semantic information in accordance with the context or behavioural goals, and acts by biasing representations in other regions (Jefferies, 2013; Lambon Ralph et al., 2017). It has been recently proposed that IFG may act differently across different phases of predictive tasks, potentially bringing together predictive coding and semantic control accounts. He and colleagues (2025) showed that connectivity between left frontal and bilateral ventral temporal cortex increased when there were strong predictions about the upcoming concept, with a subsequent reduction in connectivity if the predicted item appeared. Conversely, they reported an increase in frontal to temporal connectivity if the item could not be predicted. This points to a role for the IFG in contextually-relevant predictions of upcoming objects, and a switch to enhanced connectivity when the predicted item does not follow. This also aligns with the top-down facilitation model of object recognition, where feedback from frontal to temporal regions sends predictions about contextually relevant objects (Bar, 2004; Trapp & Bar, 2015). Within such predictive coding schemes, a mismatch between the predictions and bottom-up input will result in further enhanced recurrent connectivity.

Our research does not shed light on whether congruency effects are ultimately the result of changes to the way objects are recognized, or how objects are integrated with the scene context. Our congruency effects could also be explained through a predictive coding account of classical N400 effects observed in language (Eddine et al., 2024). A recent computational model of language comprehension can simulate N400 modulations across different conditions, where prediction error signals from the lexico-semantic layers of the model scale with the degree of unpredictability, as we see in our MEG data albeit with a potentially later latency. Whilst our analysis does not allow us to make inferences on the exact timing of the effects observed in the clusters (Maris & Oostenveld, 2007), the approximate latency might suggest they could reflect a degree of response preparation. However, we think this is unlikely. The separation between the congruent and incongruent trials seems to appear from ∼200/300 ms and continue to around 1 second. These timings, while broad, are in line with many studies showing a modulation of the N400 response to congruency across different tasks and stimuli (Coco et al., 2020; Davidson & Indefrey, 2007; Draschkow et al., 2018; Ganis & Kutas, 2003; Kumar et al., 2021; Kutas & Federmeier, 2011; Lauer et al., 2018; Mudrik et al., 2014; Truman & Mudrik, 2018; Võ & Wolfe, 2013), suggesting that what we see is a more general congruency effect rather than one driven by response preparation. A second reason we don’t think our effects relate to response preparation, is the cortical location of the effects. Our beamformer analysis localised the congruency effect to the ventral temporal cortex, which is consistent with a semantic interpretation rather than response preparation which is more commonly associated with motor, premotor and parietal cortex (Cheyne, 2013). Finally, a control analysis over motor sensors did not reveal differences between congruent and incongruent trials in either a stimulus or response-locked analysis (see supplementary figure 1).

Whilst our study does not directly reveal the nature of the representations that are modulated, our results do add critical mechanistic detail for understanding how our prior knowledge influences the ongoing processing of objects. One of our principal findings is an increase in feedback activity from the ATL to the VTC when objects are preceded by an incongruent scene. An increasing body of work acknowledges that feedback within the ventral visual pathway increases under more challenging object recognition conditions (Campo et al., 2013; Chan et al., 2011; Clarke et al., 2011; Poch et al., 2015; Rajaei et al., 2019; Schendan & Ganis, 2012; Wyatte et al., 2014). Feedback from the ATL to the VTC is linked to more demanding semantic processing of objects (Clarke et al., 2011), a shift from visual to semantic processes (Clarke et al., 2018), and linked to recognition performance (von Seth et al., 2023). TMS evidence further suggests that top-down effects from the ATL support the semantic recognition of objects (Chiou & Lambon Ralph, 2016). We argue that the modulation observed here, within the ventral visual pathway, is likely to be concerned with the semantic processing of the visual objects.

If the increase in feedback we see here has a role in the semantic processing of the object given the prior context, then this would predict that we should observe changes in the timing of when semantic representations are accessed based on congruency (Truman & Mudrik, 2018). There is abundant evidence that recognition reaction times are slower for objects that are incongruent with the scene, but this does not address the potential differences in the dynamics of when semantic representations become accessed following a congruent or incongruent scene. Recent EEG evidence does speak to this issue, where it was shown that semantic representations are seemingly evoked at similar times for all objects, both in congruent and incongruent scene contexts, but that semantic representations persisted for a longer duration when the object was incongruent with the scene (Krugliak et al., 2023). Overall, this suggests that any recurrent dynamics in the ventral visual pathway during object recognition are further increased when objects do not benefit from prior knowledge indicated by the scene.

Our results also highlight that changes in activity in the high theta and alpha frequencies reflect the relationship of the object and scene. Theta/alpha power was higher for incongruent trials compared to congruent trials, which is in line with other recent evidence showing increased power for more unexpected objects that were situated in a real-world environment (Nicholls et al., 2025). However, most research on semantic congruency effects and oscillations comes from the language domain, where it is consistently shown that theta activity increases in response to an unexpected word or a semantic violation (Bastiaansen et al., 2005, 2008; Hald et al., 2006; Packard et al., 2020; Wang et al., 2012). The effects we see here are consistent with these findings, albeit in a visual object recognition domain and with effects extending into alpha frequencies. Further, we show that these low frequency modulations localise to the ventral temporal lobe. Low frequency effects in the ventral visual pathway have been shown to reflect the higher-level visual or lexico-semantic properties of objects (Clarke, 2020; Clarke et al., 2018; Reddy et al., 2021), whilst studies across several domains have linked increases in low frequency activity with memory retrieval processes (Freunberger et al., 2008; Herweg et al., 2016; Klimesch et al., 2001). Together, this argues that the increase in low frequency activity we see reflects the additional semantic retrieval demands needed when the object recognition process does not benefit from information in the prior scene context.

Low frequency neural signals are hypothesised to reflect long-range communication between regions (Colgin, 2013; Herweg et al., 2016; Sauseng et al., 2010). Alpha activity in particular is associated with top-down signalling during visual perception (Bar et al., 2006; Chen et al., 2023; Kerkoerle et al., 2014; Mayer et al., 2016; Michalareas et al., 2016; Stecher et al., 2025), which might align with our connectivity data showing both an increase in feedback, and higher low frequency power during incongruent trials. An alternative, but not incompatible view of the low frequency modulations, is that low frequency activity decreases for objects that are congruent with the prior scene, with the decreased activity reflecting the successful access to semantic representations. Further studies linking specific frequencies to the representational content of neural signals across the conditions could seek to uncover this.

Where we are in the world clearly shapes the cognitive processes associated with recognizing an object, with our research highlighting the role low frequency oscillations play, and that top-down connectivity functions to support object processing when those objects are increasingly unexpected.

## Acknowledgments

This research was funded in whole, or in part, by the Wellcome Trust (grant no. 211200/Z/18/Z to AC) and the Isaac Newton Trust. For the purpose of open access, the author has applied a CC BY public copyright licence to any Author Accepted Manuscript version arising from this submission.

## Notes

### Competing Interest Statement

The authors have declared no competing interest.

### Summary of Updates

Revised manuscript including new analyses; Figure 5 added; Discussion section updated.

