## Supplementary figures and images for "Neural oscillations and top-down connectivity are modulated by object-scene congruency"

### supplementary figure 1

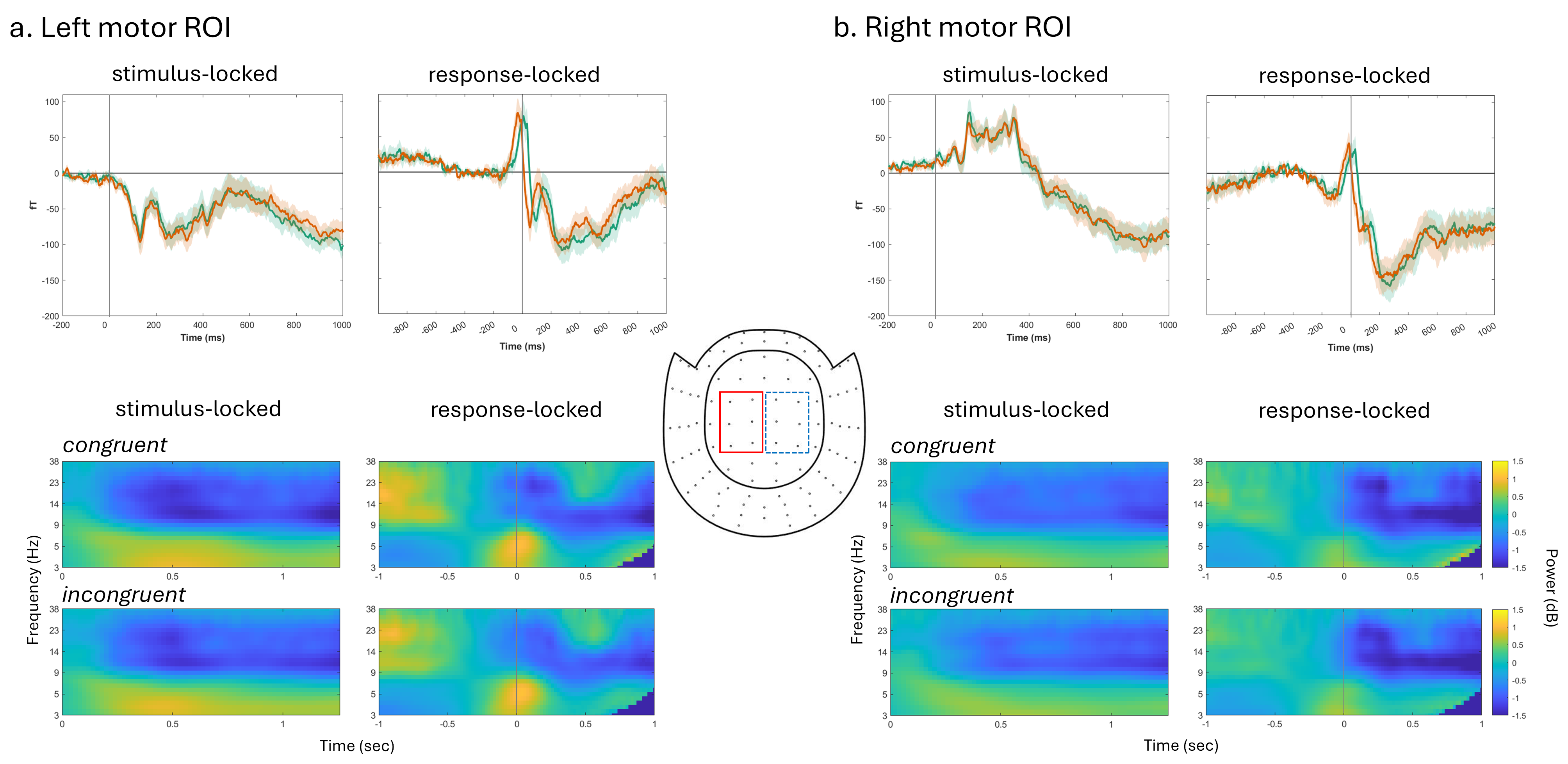
